# RIPPLE: replicate-aware detection of cell-type-anchored proximity gradients in spatial transcriptomics

**DOI:** 10.64898/2026.07.23.740288

**Authors:** Carolina Mangana, Barbara B Maier

## Abstract

Cell-cell signaling shapes tissue structure and function, yet systematically decoding these circuits with spatial transcriptomics remains an open challenge. We present RIPPLE (Replicate-Aware Inference of Paracrine Profiles via Likelihood Estimation), an R package that takes a query cell type and scans all other cell types for genes whose expression varies with distance to it. On a murine lymph node 10x Xenium dataset, RIPPLE recovers the canonical T cell zone CCL21 response program in T cells and dendritic cell subsets with unanimous sign consistency across all samples. On the public CosMx non-small cell lung cancer cohort, it identifies 515 tumor-proximity gradient genes across 17 cell types, flagging a known cancer-associated fibroblast marker (IGFBP5) as the top fibroblast hit. Overall, RIPPLE delivers ranked, cell-type-resolved paracrine candidates for experimental follow-up.

## Main

A cell’s transcriptional program is shaped not only by its own identity but also by the cells that surround it within its tissue context. Spatial transcriptomics technologies now resolve gene expression at single-cell or subcellular resolution across intact tissue sections, enabling direct measurement of how transcriptional programs differ with proximity to specific cell populations. This allows us to start addressing a central biological question: do cells near a given population (for example, tumor cells) express distinct transcriptional programs compared to cells further away? Addressing this quantitatively requires analyses that can model the relationship between expression and continuous distance while accounting for biological replication.

Existing frameworks address related but distinct questions. Spatially variable gene detection methods such as nnSVG ^1^, SpatialDE ^2^, and SPARK^3^ identify genes with non-random spatial patterns without conditioning on a cell-type anchor. STdiff ^4^ accounts for spatial correlation within a sample, but compares discrete tissue domains rather than continuous distance. Multi-view modelling approaches such as MISTy^5^ decompose expression variance into neighborhood contributions, including cell-type-specific views, and return variable-importance scores rather than directionality and a significance estimate. To complement these tools, we developed RIPPLE, an R package to provide replicate-aware, directional, cell-type-anchored gene expression gradient inference with formal statistical calibration. RIPPLE fits a per-sample Poisson generalized linear model (GLM) with a total count offset and aggregates across biological replicates via Fisher’s combined p-value under a sign-consistency gate, returning a signed gradient score and adjusted FDR for each gene in each target cell type (Fig. 1). We characterize its behavior on synthetic data, recapitulate known biology on a 10x Xenium 5k dataset, and demonstrate RIPPLE’s application to a publicly available CosMx SMI non-small cell lung cancer dataset.

**Figure 1.**
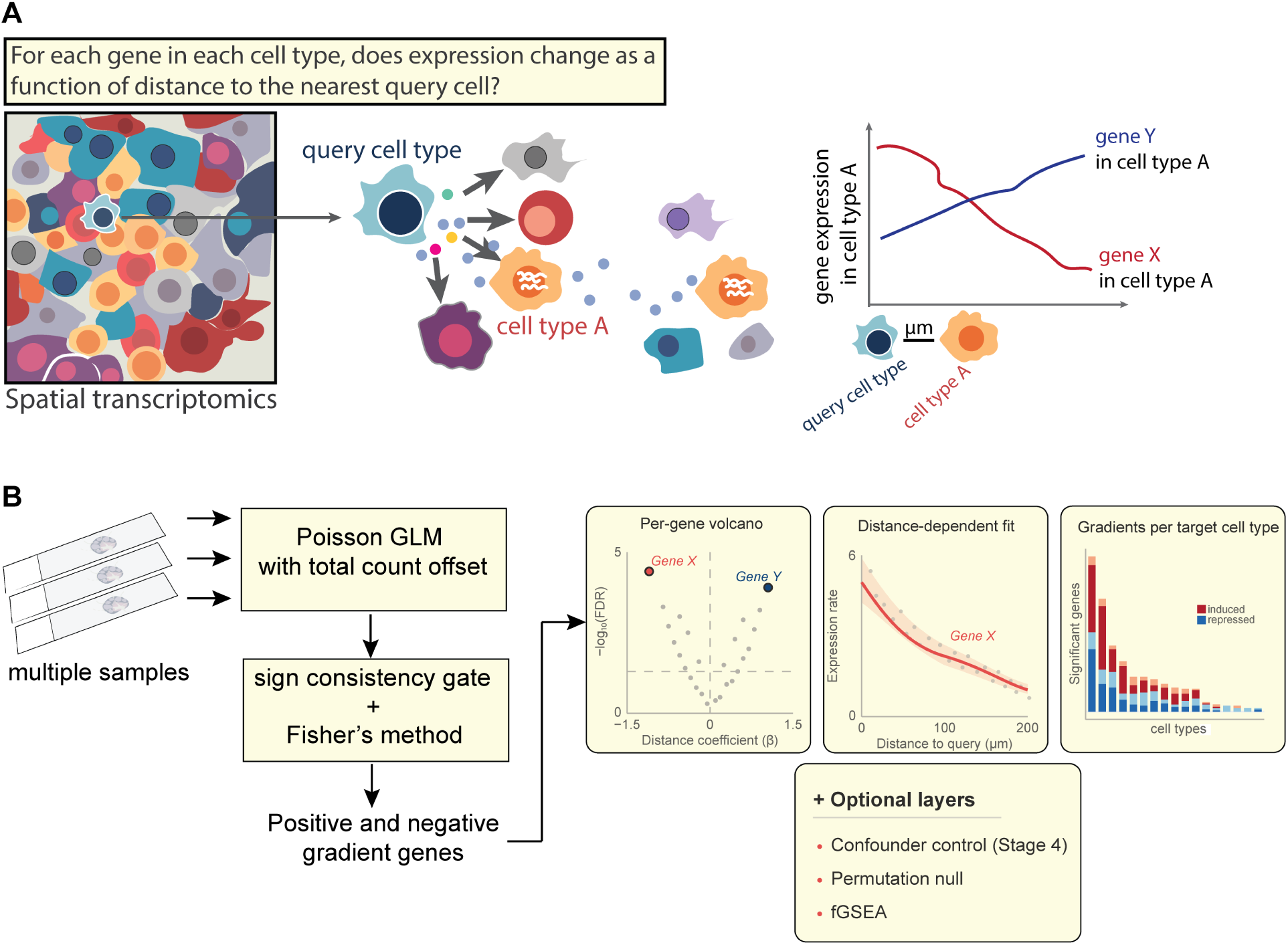
RIPPLE pipeline overview. (A) For each gene in each target cell type, RIPPLE asks whether expression rate varies with distance to the nearest query cell. The schematic shows two example genes in target cell type A: gene X has higher expression near the query (negative gradient score, induced), while gene Y has lower expression near the query (positive gradient score, repressed). (B) Per-sample workflow. From multiple biological samples, RIPPLE fits a Poisson GLM with a total-count offset for each gene in each target cell type, applies a sign-consistency gate followed by Fisher’s combined p-value across replicates, and returns the set of positive and negative gradient genes per target. The same per-gene calls feed into optional downstream layers (confounder control, permutation null, fGSEA, ligand-receptor integration), and standard outputs include per-gene volcanoes, distance-dependent gradient curves, and per-cell-type gradient counts.

### RIPPLE baseline behavior

RIPPLE was designed to test whether a gene’s expression in a target cell type changes with physical distance to a chosen query cell type (Fig. 1). It models per-cell transcript counts with a Poisson GLM whose only covariate is distance to the query cell type, so a gradient is read directly as the slope of log expression rate against distance. We further included each cell’s total counts as an offset, which turns counts into rates and prevents cells from appearing to express more of every gene simply because they were captured or segmented with more transcripts. A key strength of RIPPLE is that we fit this model separately for each biological replicate and aggregate the per-sample results into one significance value. This makes the unit of replication the sample, not the cell, enforcing that a real gradient must hold across independent samples rather than across the spatially correlated cells within one sample. These choices give RIPPLE several practical advantages. The result for each gene is a single signed number with physical units, which we call the gradient score. The sign of the gradient score gives the direction of the effect (higher or lower near the query) and its magnitude is the rate of change in expression per micrometer, which converts to a fold-change over any chosen distance. The underlying model is simple (a single Poisson regression per gene) which keeps runs fast and the assumptions easy to inspect. Because each gene is summarized by an effect size and a reproducible significance value, the output gradient lists can feed directly into downstream analyses, such as pathway enrichment and ligand-receptor inference, turning RIPPLE into a tool that can be added to existing spatial pipelines.

We first characterized RIPPLE on synthetic data designed to mimic a simplified tumor-with-lymphoid tissue (Methods). On data with no planted gradient, RIPPLE made almost no false calls. Across 75 000 null tests its pooled empirical false-positive rate (FPR) was 0.008%, 0.004%, and 0% at N = 3, 5, and 10 biological replicates (Fig. S1A). Because these null data were simulated under the same Poisson model RIPPLE fits, we also tested calibration when that assumption is broken, drawing overdispersed counts from a negative binomial (variance up to ∼2.8-fold the mean). The FPR stayed below the 5% value even at the highest overdispersion and fell toward zero as replicates increased (Fig. S1B). A per-gene ablation demonstrates how the sign-consistency gate adds a multiplicative 2·0.5^N^ safety factor on top of Fisher’s already-calibrated combination test (Fig. S1C). On data with planted gradients, RIPPLE reaches 100% power for effect sizes |β| ≥ 0.005 at every sample size (N = 3, 5, 10). For a smaller effect size (|β| = 0.002), power rose with sample size: 55%, 76%, and 85% at N = 3, 5, and 10. Relaxing the sign gate (sign_consistency = 0.75) raised power at this small effect size to 99% at N = 10 with no cost to null calibration (Fig. S1D). Finally, runtime scaled with dataset size, from 18 s on 3 000 cells to about 48 min on 250 000 cells (6.8 GB peak memory) on an Intel i7-class laptop (Fig. S1E). Real datasets with full panels in the thousands of genes and many target cell types take longer and benefit from running target cell types in parallel.

**Figure S1.**
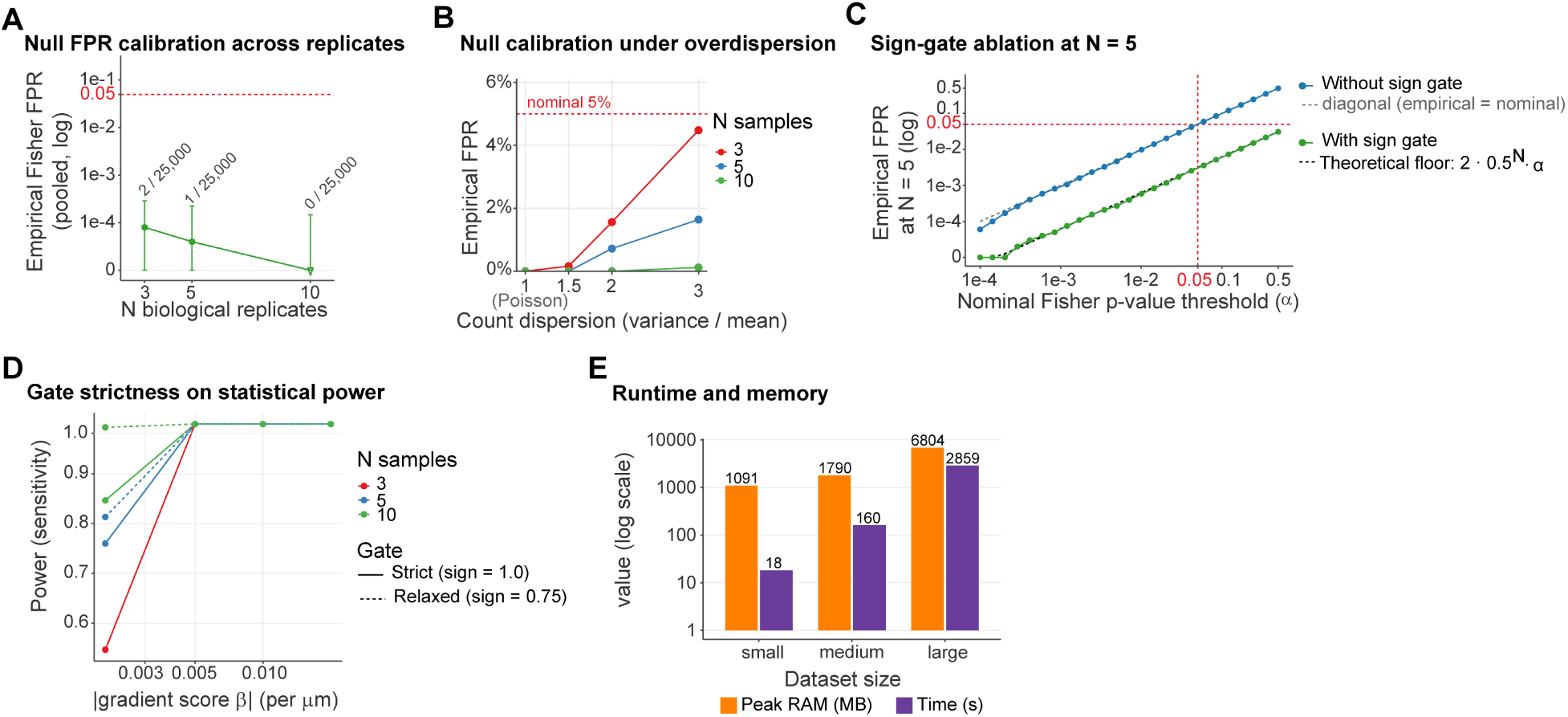
Statistical validation of RIPPLE on synthetic data. (A) Null FPR calibration. Pooled empirical false-positive rate (FPR) of the Fisher-combined test across 25 000 null tests per sample size (75 000 total across N = 3, 5, 10 replicates; 50 iterations x 500 background genes per N) on the full pipeline at default settings. Error bars: Wilson 95% CI. Downward triangle: zero false positives observed. Counts above each point: number of significant calls / total tests. (B) Null calibration under overdispersion. Empirical FPR on all-null negative-binomial data as a function of count dispersion (variance / mean; 1 = Poisson) at N = 3, 5, 10 replicates (50 iterations × 50 background genes per point). (C) Sign-gate ablation at N = 5. Combination-step simulation: 200 000 “genes”, each with N = 5 independent uniform p-values and random signs. The x-axis is the nominal Fisher p-value threshold α (log scale); the y-axis is the resulting empirical FPR. Blue: empirical FPR of Fisher’s combined p without the gate, which tracks the dashed diagonal (empirical = nominal). Green: with the gate, FPR sits on the dotted theoretical gate floor (2·0.5ᴺ·α). (D) Power to detect planted gradient-score genes as a function of effect size (|β|, gradient score per µm) and sample size, comparing strict (sign_consistency= 1.0) and relaxed (sign_consistency = 0.75) gates. Gradient score is the log-rate change per µm. (E) RIPPLE wall-clock time (seconds) and peak R-managed memory (MB) for three synthetic dataset sizes: small = 3 000 cells (1 000 per sample: 150 Tumor, 600 T, 250 Fibroblast) x 100 genes x 3 samples; medium = 50 000 cells (10 000 per sample) x 300 genes x 5 samples; large = 250 000 cells (50 000 per sample) x 500 genes x 5 samples. Values are medians across 3 replicates per size.

### RIPPLE walkthrough on a murine lymph node Xenium dataset recovers canonical T cell zone biology

Next, we set out to validate RIPPLE against known spatial biology on a real dataset and illustrate the end-to-end workflow. We ran the full pipeline on a dataset of four healthy murine cervical lymph nodes generated in-house with the 10x Xenium Mouse Prime 5K panel plus 100 custom genes (226 350 cells, companion manuscript Mangana, Neuwirth-Kodym et al. in preparation; dataset will be made available upon publication). The biological positive control question was whether RIPPLE could recover the canonical T zone program in T cells as a function of proximity to CCL21-producing T cell zone fibroblastic reticular cells (TRC). This is a continuous paracrine gradient in which TRCs secrete CCL21 to recruit and maintain CCR7+ T cells in a naive or stem-like state (characterized by high expression of Sell, Lef1, Tcf7, Il7r, Ccr7) ^6,7^.

First, we chose cell type annotation resolution. For RIPPLE, cell-type granularity directly shapes results and should be an informed choice. We recommend asymmetric resolution: the query cell type should be finely resolved to provide a biologically specific distance anchor, while target cell types can be kept broad so that gene-level expression gradients are detected rather than partitioned into subtype labels. Too fine target annotations risk circularity when the genes defining the subtypes overlap with those being tested, effectively restating the annotation as a new spatial finding (for example, if one defines a cell subtype based on the expression of a marker that determines its location). Additionally, an overly coarse query resolution dilutes cell-type-specific biology, while an overly fine query resolution can fall below the minimum cell-count threshold. Where feasible, running the analysis at two resolutions (pooled and split) provides a useful consistency check. For this analysis, we used a coarse annotation for immune and stromal identity, and a stromal re-clustering to separate fibroblastic reticular cell subtypes (TRC-Ccl21a, TRC-Cxcl12, FDC, pericytes, myofibroblasts). TRC-Ccl21a was used as the fine-grained query anchor and the eight coarse T cell subtypes (Activated_CD8, Cytotoxic_CD8, gdT_cell, Naive_CD4, Naive_CD8, Tfh, Tpex, Treg) were collapsed into a single “T_cell_all” target. We then used the built-in k-diagnostic (Fig. 2A). Mean distance to the k nearest TRC-Ccl21a cell rose smoothly for most cell types. This identified the query as a distributed population (Fig. S2) for which k = 1 is both interpretable and relatively stable. Therefore, the main analysis was run using this setting.

**Figure 2.**
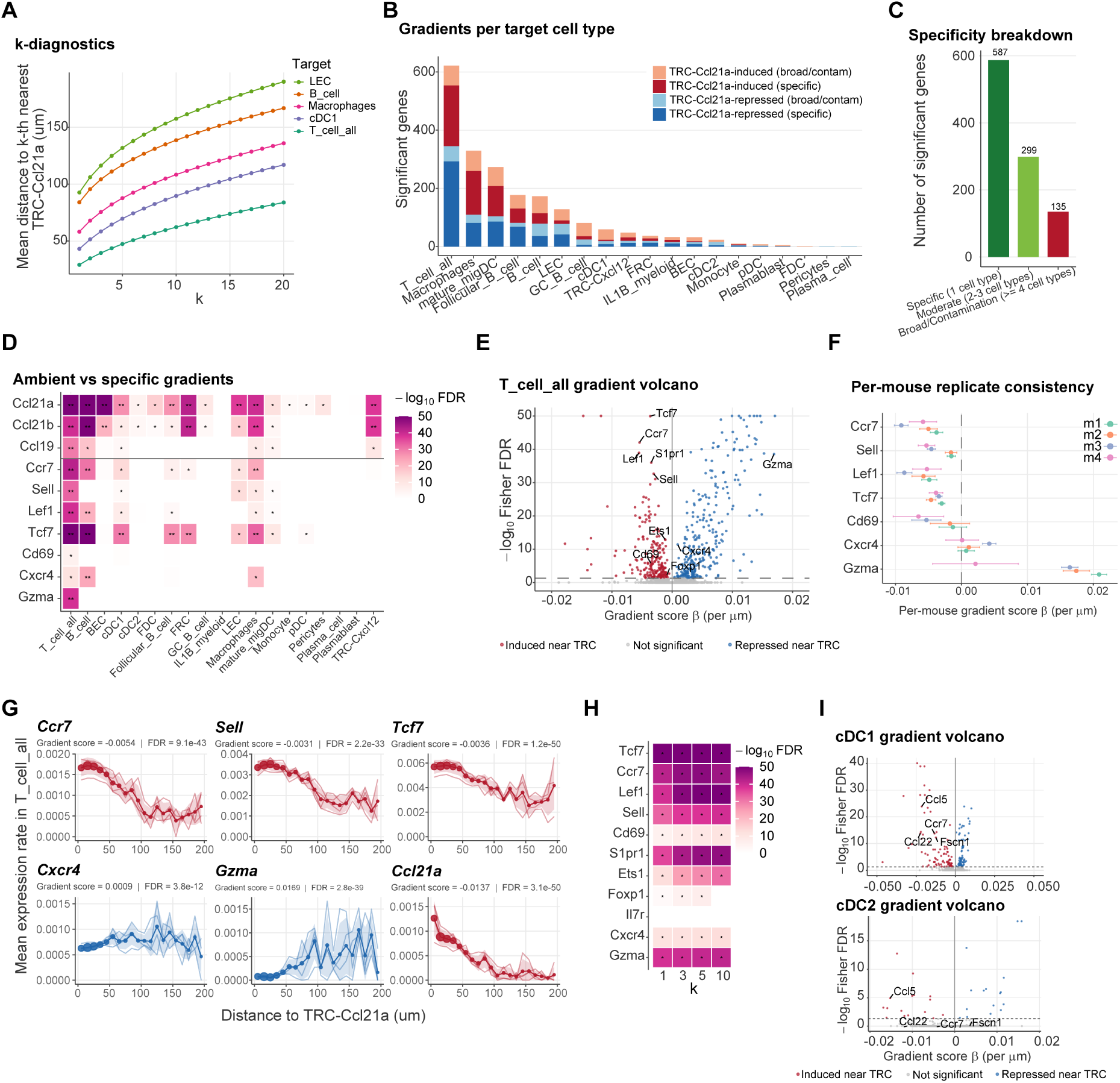
Pipeline walkthrough and validation on naive murine lymph node Xenium data. 4 mice, TRC-Ccl21a as query. (A) k-diagnostic: mean distance to the k-nearest TRC-Ccl21a cell for five representative target populations, averaged across mice. Smooth concave curves indicate a distributed query, for which k = 1 is interpretable and stable. (B) Significant gradient genes per target cell type, stacked by direction (TRC-Ccl21a-induced vs -repressed) and stratified by the cross-cell-type contamination heuristic (dark = specific; light = broad expression/potential contamination, defined as significant in ≥ 4 cell types in the same direction). T_cell_all has the most significant gradient genes. (C) Gradient gene count per target cell type across all targets, split by specificity class (specific, moderate, broad/contamination). (D) Cross-cell-type Fisher FDR heatmap (capped at −log_10_FDR = 50) for three TRC-produced chemokines (upper block) and seven canonical T zone positive-control genes (lower block). The upper block is the query-signature-on-target ambient-RNA pattern; the lower block is the cell-type-specific biology. * FDR < 0.05; ** FDR < 10^−20^. (E) T_cell_all gradient-score volcano. Red: induced near TRC. Blue: higher away from TRC/repressed. Dashed line: FDR = 0.05. (F) Per-mouse gradient score β (point) and 95% Wald CI (bar) for seven positive-control genes across the four biological replicates. (G) Binned mean per-transcript expression rate in T_cell_all as a function of distance to nearest TRC-Ccl21a. Shaded band: 1 SE across mice. Curves are red for genes with negative gradient score (induced near TRC) and blue for positive gradient score (repressed near TRC). (H) Robustness to the k_neighbors parameter. Fisher FDR heatmap of ten canonical T zone positive-control genes in the TRC-to-T_cell_all analysis at different k. * = FDR < 0.05. All positive-control gradient-score directions are preserved across k. (I) DC1 and DC2 gradient-score volcano (per-subtype run, query TRC-Ccl21a). Migratory/homing markers are labelled (Ccr7, Ccl22, Fscn1, Ccl5). Red: higher near TRC-Ccl21aquery; blue: lower expression near query. Dashed line: Fisher FDR = 0.05. In DC2, the migratory markers fall below the significance line and the gradient set is sparse. LEC = lymphatic endothelial cells; GC = germinal center; BEC = blood endothelial cells; migDC = migratory DC; TRC = T cell zone fibroblastic reticular cell; pDC = plasmacytoid DC; FRC=fibroblastic reticular cell.

Across target cell types, RIPPLE detected a wide range of significant gradient genes (Fig. 2B). T_cell_all was the top hit with ∼620 gradient genes, consistent with CCL21-driven homing being the dominant paracrine program in the T cell zone, followed by B cells, macrophages, and lymphatic endothelial cells (LEC). Gradient genes split into TRC-Ccl21a-induced (negative score; expression higher near TRC) and TRC-Ccl21a-repressed (positive score; expression higher far from TRC). They were further classified by RIPPLE’s cross-cell-type logic into specific (significant in 1 target), moderate (in 2–3 targets), or broad (in ≥4 targets) (Fig. 2C). This distinction is important for separating true signal from contamination (Fig. 2D). Before filtering, the three TRC-produced chemokines (Ccl21a, Ccl21b, Ccl19) appeared as significantly induced gradients across multiple target populations. However, this is a query-signature-on-target artefact reflecting ambient RNA or segmentation bleed-through, not real transcription in the target cell. This is mitigated by RIPPLE’s cross-cell-type logic, which flags and optionally filters any gene classified as “broad”; and the query_signature_genes argument, which lets the user name known query-produced genes (e.g. the TRC chemokines) so they are excluded from the target-cell analysis before testing. Ambient bleed-through is non-specific and therefore will show up across many cell types at once, whereas a genuine target-intrinsic gradient is restricted to the populations that truly respond. Accordingly, the seven positive-control genes (Sell, Lef1, Tcf7, Ccr7, Cd69, Cxcr4, Gzma) were mostly significant in T_cell_all, matching their known biology. Because no automated rule captures every ambient gene without false hits, combining cross-cell-type flagging with manual inspection of known query-produced ligands remains necessary. Notably, Ccr7 appears significant in multiple cell types near TRCs, but because it is broadly expressed across lymph node immune populations, this likely reflects real biology rather than contamination.

Focusing on T cells, we filtered out contamination-flagged genes. Then, we plotted the remaining genes as a gradient volcano (Fig. 2E). The canonical positive-control T cell zone program dominates the top hits. Four of five expected induced markers reached significance with unanimous sign consistency across all biological replicates: Tcf7, Ccr7, Lef1, and Sell. Il7r showed a coefficient in the expected direction but failed the sign-consistency gate (0.5). Cd69, a T cell retention marker expected to be highest where cells are held in place by chemokine signaling^8^, was also significantly induced near TRC-Ccl21a. In contrast, Gzma, a cytotoxic granzyme expected to be repressed in the naive T cells^9^ which dominate the T cell zone, was significantly higher expressed far from TRC-Ccl21a, consistent with effector-program cells localizing away from the naive T cell zone^10^. Cxcr4, which can guide cells toward Cxcl12+ fibroblasts, was similarly increased away from TRC-Ccl21a, likely mirroring the known spatial segregation of Cxcl12+ from Ccl21+ fibroblasts in lymph nodes^11^. This highlights how RIPPLE outputs both direct effects and consequences of tissue structure. Using another of RIPPLE’s visualization functions, we can inspect replicate-level variation (Fig. 2F) to confirm that the aggregate significance is not driven by one or two outlier samples. The positive-control genes each showed the same coefficient sign in all mice, with overlapping 95% Wald confidence intervals and similar effect magnitudes. Built-in binned distance-dependent gradient curves (Fig. 2G) can be used to further visualize the continuous-gradient of the effect: Ccr7, Sell, and Tcf7 mean expression rates rise toward the query; Cxcr4 and Gzma fall toward the query; and Ccl21a itself showed the expected decay from high at zero distance, since it is a query-produced ambient ligand.

To test how sensitive the positive-control result is to the choice of k parameter, we re-ran the TRC-Ccl21a to T_cell_all analysis at different k (1,3,5,10), keeping all other parameters at their defaults (Fig. 2H). Night of eleven canonical T zone genes retained their expected coefficient direction at every k with unanimous sign consistency across the four biological replicates. Fisher FDR either improved with larger k or remained the same. Coefficient magnitudes were stable in sign and order of magnitude. For example, Ccr7 varied from -0.0054 to -0.0060 and Gzma from +0.0169 to +0.0173 across the four k values, validating the choice of k we made based on the k diagnostic plot. The interpretation of the analysis is therefore unchanged across the tested range. Nonetheless, two edge cases appear. Foxp1, a low-abundance transcription factor (pct_expr ∼0.38), was significant at low k but dropped below the sign-consistency gate at k=10 (consistency fell to 0.75), demonstrating that averaging over long neighborhoods can destabilize genes with sparse per-sample signal. Il7r remained non-significant at every k, likely due to difficulty in capturing any transcripts, showing that this reflected genuine per-replicate heterogeneity in this dataset. Overall, these observations are evidence that for a distributed query cell such as TRC-Ccl21a, k acts as a smoothing dial that recovers significance without altering direction or sign, but for isolated low-expression genes, higher k can bring the per-replicate consistency near the gating threshold. This may be particularly important for cytokine signals, which are often lowly expressed. As a second test of T cell zone biology, we asked whether RIPPLE could recover the CCR7-dependent retention of type I dendritic cells (DC1) in the deep T cell zone in homeostasis.

Dendritic-cell subtypes occupy distinct niches in the lymph node at homeostasis, with DC1 populating the parenchyma and T cell zone, and DC2 localizing to the interfollicular areas and T-B border zone^12,13^. Accordingly, only DC1 should express Ccr7 as a steep gradient responding to the TRC-Ccl21a signal. The DC1 gradient gene set had mostly induced markers (Fig. 2B, Fig. 2I), and Ccr7 was recovered as a strongly induced gradient (median gradient score = −0.013, Fisher FDR = 1.8 × 10⁻¹⁴, Fig. 2I). Therefore, as expected, DC1 express progressively more CCR7 the closer they sit to the CCL21-producing T-zone stroma. The migratory-DC chemokine Ccl22 showed the same DC1-restricted induction. In DC2, the same genes did not exhibit gradients, the volcano plot is sparse (Fig. 2I), and Ccr7 showed only a weak, non-significant distance profile. DC1 and DC2 had comparable numbers of genes pass the expression filter (1 772 vs 1 756), showing that this contrast is not a power difference. Nevertheless, Ccr7’s DC1 coefficient is about four times its DC2 value. DC2 did return fewer significant gradients overall (32 vs 173), so here Ccr7 is a weak, non-significant gradient rather than a strict absence. This reflects the spatial biology of homeostatic lymph nodes, wherein DC1 are attracted by Ccl21-producing TRCs into the deep T cell zone, whereas DC2 respond to additional signals that control their distinct positioning.

Overall, RIPPLE can be successfully used to recover and visualize known chemokine-induced gradients in immune cells, which can be extended to uncover new spatial biology.

### Applying RIPPLE to public CosMx non-small cell lung cancer dataset

We also applied RIPPLE to the public CosMx SMI non-small cell lung cancer dataset from He et al. ^14^. This dataset comprises more than 750 000 cells across 5 patients profiled with a 980-gene panel. Patient-specific tumor subtype labels were merged into a single “tumor” annotation, and this population was used as the query. The full pipeline was run with default parameters (Fig. 3A, k = 1 nearest neighbor, max distance 200 µm, Fisher FDR threshold 0.05) using patient as the biological replicate unit. RIPPLE identified 515 distance-dependent gene gradients across 17 target cell types (Fig. 3B). Fibroblasts showed the most extensive tumor-proximity remodeling, followed by macrophages, CD4 memory T cells, endothelial cells, neutrophils, and mDCs (mature DCs). The top-ranked fibroblast gene by Fisher FDR was MIF. However, RIPPLE’s cross-cell-type logic flagged it as a likely contaminant because it was significantly induced in 9 out of 17 target cell types, matching an ambient RNA or segmentation profile rather than fibroblast-specific expression. After filtering broadly modulated genes, the top fibroblast hit was IGFBP5 (*β* = -0.0037, Fisher FDR = 2 × 10^−5^^1^, induced near tumor in all 5 patients; Fig. 3C, D). Notably, this encodes a protein known to be often expressed in cancer-associated fibroblast activation^15^. Other fibroblast hits are matrix and stromal genes that are higher in tumor-distal fibroblasts: CXCL12, MMP2, THBS1, and PTGDS, which are all known cancer-associated fibroblast genes^16–18^. This expression pattern reflects cancer-associated fibroblast spatial distribution where activated CXCL12+ fibroblasts, main mediators of immune exclusion^19^, are not positioned immediately next to cancer cells but rather ∼100µm away, defining an architecture for stromal activation.

**Figure 3.**
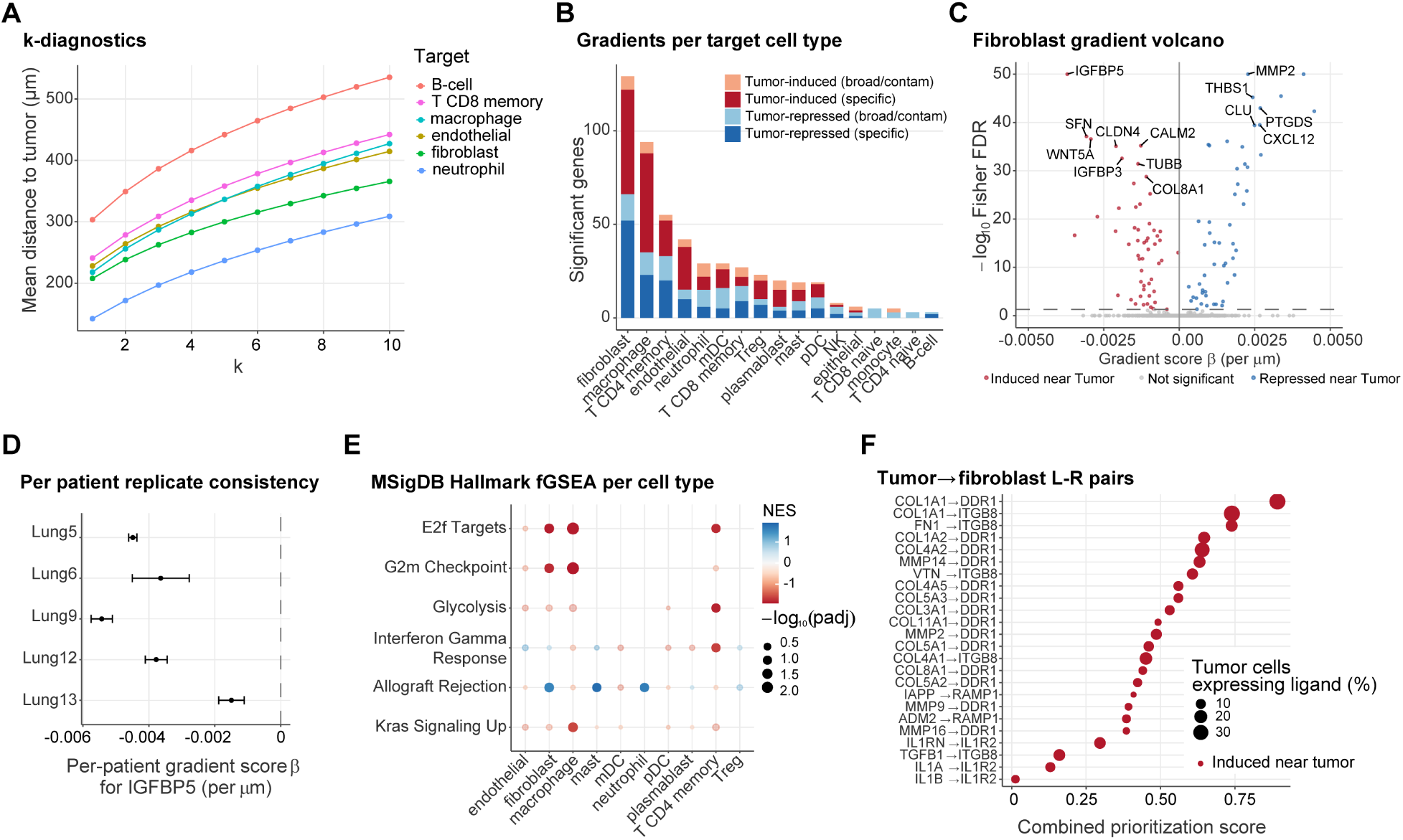
RIPPLE applied to the CosMx NSCLC dataset. 5 patients, tumor cells as query. (A) Distance from each target cell type to the nearest tumor cell as a function of k, per patient. (B) Number of significant gradient genes per target cell type. (C) Fibroblast gradient volcano. IGFBP5 highlighted. (D) Per-patient forest plot of the IGFBP5 gradient score. (E) MSigDB Hallmark fGSEA across target cell types. Color = NES, dot size = −log₁₀(padj). Only pathways at padj < 0.1 are shown. Cell types with fewer than 100 ranked genes were not tested and cell types with no pathway reaching significance are omitted. (F) Top ligand–receptor pairs from the fibroblast L-R integration layer.

RIPPLE gradient lists are signed and thus can be used as input for several existing tools. Built-in MSigDB Hallmark fGSEA integration of RIPPLE results ranked by gradient score (Fig. 3E) showed proliferation programs induced near tumor: E2F targets in macrophages, fibroblasts, and CD4 memory T cells, and G2M checkpoint in macrophages and fibroblasts. Macrophages near tumor also showed KRAS signaling, and CD4 memory T cells showed glycolysis and an interferon-gamma response. The only program enriched away from tumor was allograft rejection, repressed near tumor across fibroblasts, mast cells, and neutrophils.

Genes marked as induced by RIPPLE can also be interpreted as a gene set of response to the query cell. This allows us to explore upstream inducers of that response to help decipher potential signaling mechanisms between the query cell and the target cell. To this end, we ran a ligand-receptor integration layer on fibroblasts as the receiver, and ship this as an optional built-in analysis step. In summary, we took the fibroblast gradient hits, filtered them to known receptors from the NicheNet v2^20^ curated ligand-receptor network and, for each candidate receptor, scored the set of tumor-expressed ligands that can plausibly signal through it using three lines of evidence. The first is a direct score (the fraction of tumor cells expressing the ligand times the absolute gradient coefficient of its receptor on fibroblasts), capturing pairs where a well-expressed tumor ligand has a spatially modulated receptor on the receiver. The second is NicheNet’s pre-computed ligand-activity AUROC^20^, which asks whether the full fibroblast gradient-gene set is enriched in that ligand’s known transcriptional targets. The third is a downstream target enrichment test that checks whether the fibroblast gradient genes are themselves enriched for the ligand’s direct targets. These three scores, plus the ligand’s overall tumor-expression prevalence, are each rescaled across candidate ligand-receptor pairs and combined into a weighted average. The top tumor to fibroblast pairs after broad-expression filtering (Fig. 3F) are dominated by ECM remodeling and matrix-receptor biology, for example, collagen to DDR1 pairings (COL1A1 → DDR1, COL1A2 → DDR1) and matrix-metalloproteinase signaling (MMP14 → DDR1, MMP2 → DDR1). Matrix proteins also engage integrin ITGB8 (FN1 → ITGB8, VTN → ITGB8), and two signaling families: TGF-β (TGFB1 → ITGB8) and interleukin-1, in which the tumor IL-1 ligands and the IL-1 antagonist converge on the decoy receptor IL1R2 (IL1A, IL1B, IL1RN → IL1R2). This pattern of inflammatory signaling and collagens and matrix-remodeling enzymes engaging DDR1 and integrin receptors on adjacent fibroblasts is consistent with previously described cancer associated fibroblast activation^21^. Thus, RIPPLE faithfully recapitulates known cancer-associated fibroblast signaling and can be integrated with L-R analysis tools to turn its signed gradients into a ranked shortlist of candidate interactions for experimental follow-up.

Overall, RIPPLE can be applied to public datasets with minimal effort, allowing for the discovery and prioritization of targets.

## Discussion

RIPPLE fills a specific niche in the spatial transcriptomics toolkit by providing replicate-aware, directional, cell-type-anchored gene expression gradient inference. Its main design choices, per-sample fitting followed by Fisher aggregation and a sign-consistency gate, address pseudoreplication by treating biological replicates, not cells, as the unit of inference and requiring gradients to agree in direction across independent samples. The sign-consistency gate intentionally trades power for reproducibility. We recommend the default of 1.0 and propose relaxing to 0.75 only with at least 6 replicates where one discordant sample is biologically plausible.

Other recent methods address problems related to those we try to tackle here. Ospina et al.^4^ proposed STdiff, which incorporates spatial covariance structures into linear mixed models. RIPPLE addresses the same underlying concerns but at a different level: rather than modelling spatial covariance within a sample, it aggregates across biological replicates. The two approaches answer complementary questions. For users concerned about residual within-sample spatial autocorrelation in their RIPPLE fits, the check_spatial_autocorrelation() function provides a post-hoc diagnostic.

Several limitations of RIPPLE should be kept in mind when interpreting results. First, like all current spatial transcriptomics tools, RIPPLE is transcript-based: a significant gradient means a gene’s RNA abundance per cell varies with distance to the query cell type, not that the query directly secretes a factor acting on the target. Mechanisms such as cytokines, ligands, or contact-mediated signals must be inferred and require orthogonal validation (protein-level assays, perturbation experiments). Second, the optional confounder control (Stage 4) fits only a single additional distance covariate in a bivariate GLM and does not capture non-linear effects, covariate interactions, or unmeasured confounders. Thus, its classifications (query_specific, enhanced, niche_driven) should be treated as hypothesis-generating. Additionally, one can re-run RIPPLE on a cell type that shares the niche with our initial query and evaluate how much signal is still recovered there. Finally, RIPPLE supports only a single query cell type as the distance anchor and detects transcriptional gradients, not co-localization: a gene whose positive cells cluster spatially but whose per-cell expression does not vary with distance will not be flagged. Therefore, we recommend using RIPPLE alongside complementary co-localization analyses.

Overall, RIPPLE is an additional tool for single-cell spatial transcriptomics. By measuring how expression changes with distance rather than simple co-localization of two cell types, and by requiring that gradient to remain consistent across biological replicates, RIPPLE generates testable hypotheses about how cells influence one another within complex tissues.

## Methods

### Statistical model

For each gene in each target cell type in each biological replicate, RIPPLE fits a Poisson GLM to the per-cell raw transcript counts:

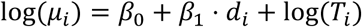

where *μ_i_* is the expected count for cell *i*,

*d_i_* is the Euclidean distance (in µm) to the nearest query cell (k = 1 nearest neighbor, or user defined k number), and *T_i_* is the total transcript count per cell *i*, included as an offset to convert raw counts to rates. Cells further than 200 µm from the nearest query cell are excluded from the fit by default because most paracrine signaling acts within this range. The user can adjust the cutoff by changing the function argument for maximum distance, if needed to match the biological question and tissue architecture. The coefficient *β*_1_ is the log-rate change per micrometer. We refer to this per-gene per-sample coefficient *β*_1_ as the gradient score and use this term interchangeably with *β* in the Results, figure captions, and axis labels throughout the paper. The gradient score is signed, with negative meaning higher expression near the query (induced) and positive meaning the expression is lower near the query (repressed).

The Poisson model assumes the mean equals the variance. While modest overdispersion is tolerated in practice, it can be checked. RIPPLE writes a per-cell-type dispersion summary as part of every run, and users should inspect this output: if overdispersion is a concern for a certain dataset and the Poisson assumption is violated, a quasi-Poisson or negative binomial GLM would be more appropriate. In practice, for current high-plex imaging panels (Xenium 5K, CosMx), where most genes per cell are sparsely detected and the count distribution is approximately quasi-binary, the Poisson default performs well. We have not observed median dispersion approaching >2 in either dataset.

### k-neighbors selection and diagnostics

The k_neighbors parameter controls how “distance to the query cell type” is defined. At *k* = 1 the predictor is the Euclidean distance to the single nearest query cell, whereas at *k* > 1 it is the mean distance to the *k* nearest query cells. Neither choice is universally applicable, and the right value depends on the spatial distribution of the query population (Fig. S2). The chosen k should also be considered for interpretability of the gradient score.

At *k* = 1, the predictor is easily interpretable (how far is the closest query cell?), but it is vulnerable to several pitfalls. A single mis-segmented or mis-labelled query cell dominates the distance estimate for all target cells in its neighborhood, so annotation error rates can more noticeably perturb results. Additionally, segmentation over-splitting, common on imaging-based platforms, creates duplicated query fragments at near-identical coordinates that drag target-to-query distances toward zero artefactually. When the query population is clumped (for example follicular dendritic cells in lymph node follicles, tumor cells in nests, neutrophil aggregates), *k* = 1 distance is small everywhere while query is inside the clump and jumps sharply at the boundary. The linear Poisson term fits this near-step with a single slope, so most of the model’s resolution is absorbed by the boundary transition rather than by within-clump or smooth-distance variation. *k* = 1 also ignores variation in local query density: two target cells at the same distance to their nearest query may sit at the edges of very different-sized clumps.

At *k* > 1, the above artefacts are partially averaged away but new interpretability problems arise. “Mean distance to the five nearest query cells” conflates distance-to-cluster with within-cluster density, and the resulting coefficient *β*_1_ describes a log-rate change per unit of this composite rather than per micrometer of a clearly defined goal. Higher *k* can also smooth over real biological structures and tissue architectures (e.g. tumor boundary), potentially transforming a step into a gradient that is an artefact of averaging rather than an inherent property of the biology at hand. Finally, when the density of query cells varies across biological replicates (for example, different sample sectioning accuracy or plane orientation), *k* > 1 distances can differ systematically across mice in a way that *k* = 1 does not, which can produce Fisher-consistent gradients that reflect technical replicate differences in query abundance rather than shared biology. For the user, the implication is that *k* selection should be considered rather a default. RIPPLE provides plot_k_diagnostics(), which computes per-cell mean distance to the *k* nearest query cells across k (1-10) and reports how the mean and standard deviation of that distance evolve. The mean-distance curve is the primary diagnostic: a smooth, concave rise with no plateau or sharp discontinuity indicates a well-distributed query population for which *k* = 1 is interpretable and stable (as in Fig. 2A), whereas an early plateau or a step-like discontinuity indicates a clumped or sparse query whose *k* = 1 distances are dominated by clump boundaries. The standard-deviation curve is a complementary check: a sharp drop as k increases means the k = 1 estimate is noisy and a larger *k* stabilizes it, while a gradually changing SD means k = 1 is already stable. For a clumped or sparse query the user should apply a larger k, or run two resolutions in parallel and assess stability of genes of interest across *k* values. We recommend running a k-sweep and verifying that positive-control genes remain significant with the expected sign across k values.

**Figure S2.**
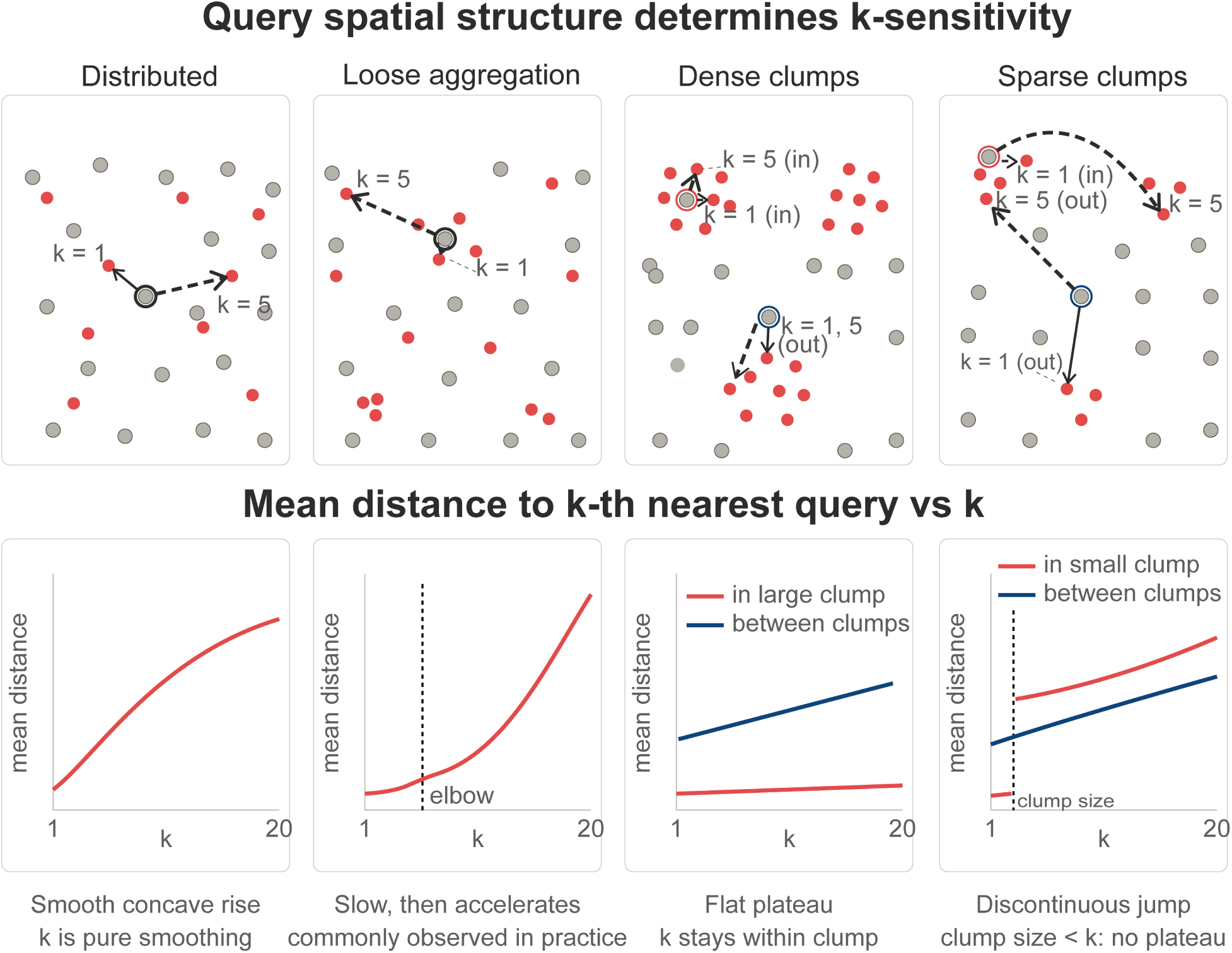
Query spatial structure determines k-sensitivity. (A) Illustrative tissue layouts for four hypothetical query distributions showing the relationship between k = 1 and k = 5, neighbor arrows from a representative target cell. (B) Schematic mean-distance-to-k-nearest curves per scenario. The plateau or its absence in the per-sample diagnostic (plot_k_diagnostics) helps the user pick a k.

### Cross-replicate inference

A sign-consistency gate is applied before the combined statistic is computed. By default, the gate requires all samples to agree on the sign of the coefficient *β*_1_. Genes failing the gate are assigned a combined p-value of 1 and are not reported as significant regardless of individual p-values. This matters most at small sample sizes. For large sample sizes, the threshold is configurable (ripple_config option sign_consistency, default is 1.0; a relaxed value of 0.75 is only suggested for designs with at least 6 replicates). Users should note any relaxation of the gate explicitly. Relaxed-gate results are less reliable than the default.

Per-sample *s*, coefficients *β*_1,*s*_ and their p-values from Wald z-tests are combined across biological replicates using Fisher’s method of combined probability test^22^. The combined p-value is computed in base R as the upper tail probability via stats::pchisq with df = 2*S*. Fisher’s combination is best when no prior weight is placed on individual replicates. This matches RIPPLE’s design choice of equal weight per biological replicate. A gene must pass the expression filter in at least two valid samples for a combined p-value to be computed; genes with fewer are reported as untested. FDR is controlled by the Benjamini-Hochberg method applied to the combined p-values within each target cell type, treating each target as a separate family of tests. Whenever we are talking about the **gradient score** for a gene across samples, we are referring to the median of individual per-sample coefficients *β*_1,*s*_ (equal weight per replicate) and the significance derived from the Fisher combination.

Finally, RIPPLE does not require every replicate to be individually significant. A hard majority rule would drop genuine gradients that only reach per-sample significance in some replicates, especially in samples with few cells. Instead, RIPPLE reports n_sig_samples, the number of replicates significant at 0.05. An optional gate (min_sig_fraction, default 0, off) enforces a minimum fraction for users who want to apply this stricter threshold.

### Expression filtering

Genes are filtered before model fitting using a two-tier system. Regular genes must be expressed in at least max(1% of cells per sample, 25 cells) in all valid samples (strict filter). A user-defined list of priority genes (for example, cytokines and receptors) is subject to a lenient filter requiring only 25 expressing cells in at least x samples (x default is 2, but should be customized). This can rescue sparsely expressed but biologically important transcripts such as cytokines, which is often worth doing for sparse spatial transcriptomics data. Samples with fewer than 30 cells of the target type are excluded from that target’s analysis.

### Cell-size offset

The offset(log(total_counts)) term converts raw count modelling to rate modelling and is critical. Cells near the query cell type may appear to express more of every gene due to ambient RNA contamination or segmentation artefacts that co-vary with cell size. Without the offset, such artefacts would manifest as negative coefficients across many genes simultaneously. With the offset, the model tests whether expression rates (not raw counts) change with distance.

### Optional confounder control

When a second cell type co-localizes with the query cell type (for example, cancer-associated fibroblasts co-localizing with tumor cells), RIPPLE can fit a bivariate model:

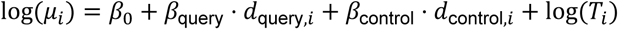

This way genes significant in Stage 1 (RIPPLE run for query) are then reclassified based on the behavior of *β*_query_ after controlling for *β*_control_ as: query_specific when Fisher FDR < threshold in bivariate model, same sign as Stage 1; enhanced when query_specific and |*β*_query_| is increased by at least 10% vs Stage 1; niche_driven when Fisher FDR is above threshold and |*β*_query_| is attenuated by at least 50%; reversed when Fisher FDR < threshold but the sign flips relative to Stage 1; and unresolved for any gene not matching the above, reported without a confident label. Note that this approach only fits a single additional distance covariate in a bivariate GLM and does not capture non-linear effects or other confounders. Hence, these classifications should be treated as hypothesis-generating only.

### Permutation validation

To validate query specificity, RIPPLE supports location-based permutation via a GPU-accelerated PyTorch script (inst/python/run_permutation_gpu.py) or a pure-R fallback. For each permutation, within each sample, N cells are drawn at random from the full cell pool (query, target, and other cell types), where N equals the original query cell count in that sample. These pseudo-query cells become the new distance anchor, target-to-pseudo-query distances are recomputed, and the per-sample Poisson GLM is re-fit to generate an empirical null distribution of coefficients. The procedure preserves the per-sample query count and the spatial distribution of cells but randomizes which cells serve as the distance anchor. For each permutation, the per-sample null coefficients are summarized by their median, the same statistic as the reported gradient score, and an empirical p-value is obtained by comparing the observed median coefficient to this null distribution. The Fisher combination and sign-consistency gate used for the observed analysis are not applied on the permutation path, so this step checks query-location specificity of the effect size rather than re-deriving the reported significance.

Two design choices deserve attention. First, because pseudo-query cells are drawn from the full cell pool, including the target population, a target cell can by chance be selected as a pseudo-query, giving it a self-distance of zero and shifting the null distribution toward smaller distances. This makes the permutation conservative for abundant target populations, where self-selection is common. Second, the null tested is location specificity of the query, not specificity against a particular alternative cell type. If the biological question is whether a gradient is driven by the query rather than by another cell type that happens to co-occupy similar spatial niches, a targeted permutation restricted to the two populations of interest is more informative than the default. If the question is the existence of any spatial structure rather than query specificity, an SVG method (nnSVG, SpatialDE) is more appropriate. The permutation design should therefore be selected to match the biological question and be interpreted carefully.

### Spatial autocorrelation diagnostic

We also provide an additional check_spatial_autocorrelation() function that computes Moran’s I on deviance residuals for a user-supplied set of genes, using spdep::moran.test() with a k-nearest-neighbor spatial weights matrix (k = 20, default). Residuals with |*I*| < 0.05 are considered consistent with the independence assumption of the GLM; values above 0.15 indicate non-negligible residual autocorrelation, so the within-sample Wald p-values should be interpreted with caution; per-sample fitting and Fisher aggregation across replicates partially mitigate this, and permutation-based significance can serve as a cross-check. This is not run by default. We recommend it for advanced users when residual within-sample autocorrelation could affect specific genes of interest.

### Pathway enrichment

The run_ripple_fgsea() function ranks genes within each target cell type by their gradient score *β*_1_ and runs fast preranked gene-set enrichment analysis ^23^ against MSigDB collections (Hallmark by default; KEGG, Reactome, and GO:BP also supported). Because the gradient score is signed, fGSEA returns directional normalized enrichment scores: positive NES indicates pathway genes are concentrated among the most repressed-near-query genes, negative NES indicates concentration among the most induced-near-query genes. Because spatial transcriptomics panels are small (typically a few hundred to a few thousand genes), large MSigDB collections (Hallmark, Reactome; 50–200 genes per set) overlap the panel only sparsely and can yield under-powered enrichment statistics. For these settings the user can also supply compact custom curated modules (e.g. 5–15 genes per set) through the gene_sets argument of run_ripple_fgsea().

Pathway enrichment results should be examined gene by gene before any biological claim is made. The aggregate NES can be driven by a small number of strongly gradient-associated genes, by idiosyncratic gene-set composition, or by contamination-flagged genes that still pass the default expression filter if the user does not select the argument to remove them. We recommend three sanity checks for any leading pathway:

1. inspect the leading-edge gene set explicitly;
2. cross-reference it against the contamination candidate list emitted by the cross-cell-type flagging heuristic or run fGSEA only on non-contamination candidates;
3. verify that the per-sample gradient score direction is consistent across replicates for each leading-edge gene. The package provides gradient-curve and per-replicate visualization helpers (plot_gradient_curve(), create_forest_plot()) that support this gene-level inspection without leaving the pipeline.

### Synthetic data design

For testing RIPPLE on synthetic data, SpatialExperiment objects were generated to imitate a simple tumor structure. Each dataset contained three cell types: a Tumor cluster of 30 cells sampled uniformly from a disc of radius 60 µm at the center of a 500 µm field, 120 T cells sampled uniformly across the field, and 50 Fibroblasts sampled from a ring of inner radius 100 µm and outer radius 200 µm centered on the tumor cluster. Per-cell total counts were drawn from a Poisson distribution with mean *b* ⋅ *G* (where *b* = 3 is the background rate and *G* is the number of genes), and a size factor was derived as the ratio to the expected total. For null datasets, per-gene per-cell rates were set to *b* multiplied by the size factor. For gradient datasets, a specified number of genes in T cells only had their rates multiplied by exp(β·d) for negatively biased genes (induced near tumor) or exp(−β·d) for positively biased genes (repressed near tumor). Here β is signed (default −0.01); for induction near the tumor the exponent decreases with distance, so induced genes carry a negative gradient. The Poisson null calibration (Fig. S1A) used 500 background genes with no planted gradient across 50 iterations per sample size (25,000 null tests per N), and the power benchmark (Fig. S1D) used 5 planted gradient genes with 45 background genes across 30 iterations at four effect sizes, comparing the strict (1.0) and relaxed (0.75) sign-consistency gates. Both used 200 cells per sample (30 Tumor, 120 T, 50 Fibroblast). The sign-gate ablation (Fig. S1C) is a separate p-value-level simulation and does not use the synthetic tissue. All simulations used fixed per-benchmark base seeds and deterministic per-iteration offsets to ensure reproducibility across the benchmarks. To test calibration when the Poisson mean-equals-variance assumption is violated, we additionally generated all-null datasets in which per-cell counts were drawn from a negative binomial with a constant variance-to-mean ratio (1.5, 2, 3; 1 recovers the Poisson case), using 50 background genes across 50 iterations, while still fitting the standard Poisson GLM (Fig. S1B).

## Software and data availability

### Software

RIPPLE is implemented as an R package available at https://github.com/Maier-Lab/RIPPLE. Five vignettes and an interactive webpage ship with the package: a getting-started guide, a walkthrough on the CosMx NSCLC dataset, a comparison with alternative spatial tools, a guide on how to run parallel RIPPLE runs for larger datasets, and a rendering of the FPR calibration and power benchmarks. The interactive page explains RIPPLE’s core design principles and is available at https://maier-lab.github.io/RIPPLE/.

The package requires R ≥ 4.0 and pulls core dependencies (Seurat^24^, data.table, ggplot2^25^, patchwork^26^, RANN^27^, ggrepel^28^, Matrix^29^, scales, pheatmap^30^, spdep^31^) from CRAN. Optional functionality requires additional Bioconductor packages (fgsea^23^, msigdbr^32^, SpatialExperiment^33^) and NicheNet^20^ (from GitHub), listed in Suggests. GPU-accelerated permutation testing is provided via a bundled Python script using PyTorch.

For runs on full panels (e.g. Xenium 5K) with many target cell types, we recommend running RIPPLE in parallel across target cell types whenever possible, since each target cell type is fit independently. We provide a vignette to guide users through parallelization. A single-core run sweeps the target cell types in series and can take many hours on dense panels; parallelizing the sweep with future^34^ or splitting it into one job per target cell type on a SLURM cluster drops the wall-clock to the slowest target cell type. SLURM array job templates for this pattern are bundled in inst/slurm/ of the RIPPLE repository. As a concrete reference point, a single target cell type with 79 043 B cells across 4 mice on a Xenium 5K panel (2 412 genes passing the expression filter) ran in approximately 29 minutes on a 4-CPU SLURM node.

### Data

The CosMx SMI non-small cell lung cancer dataset analyzed in this manuscript was published by He et al.^14^ and is available from the original authors’ release. The naive murine cervical lymph node Xenium dataset is from a companion manuscript currently in preparation (Mangana, Neuwirth-Kodym et al.); raw data will be released with that manuscript. All synthetic benchmark datasets are reproducibly regenerated from seeded scripts in data-raw/benchmarks/ of the RIPPLE repository.

## Acknowledgements

Thank you to Raphael Bednarsky, David Fisher, and Florian H. Schmidt for discussions on RIPPLE parameters and functions. To Teresa Neuwirth-Kodym for writing input. To Mariia Naumova and Sarah Dobner for testing and user experience feedback.

